# Immunogenicity of autologous and allogeneic human primary cholangiocyte organoids

**DOI:** 10.1101/2024.01.11.574744

**Authors:** Sandra Petrus-Reurer, Olivia Tysoe, Winnie Lei, Maelle Mairesse, Thomas Tan, Sylvia Rehakova, Krishnaa Mahbubani, Julia Jones, Cara Brodie, Namshik Han, Catherine Betts, Ludovic Vallier, Kourosh Saeb-Parsy

## Abstract

Primary human cells cultured in 3D organoid format have great promise as potential regenerative cellular therapies, but their immunogenicity has not yet been fully characterized. In this study, we use *in vitro* co-cultures and *in vivo* humanized mouse experimental models to examine the human immune response to autologous and allogeneic primary cholangiocyte organoids (PCOs). Our data demonstrate that PCOs upregulate the expression of HLA-I and HLA-II in inflammatory conditions. The immune response to allogeneic PCOs is driven by both HLA-I and HLA-II and is substantially ameliorated by donor-recipient HLA matching. Autologous PCOs induce a low-level immune infiltration into the graft site, while allogeneic cells display evolving stages of immune rejection *in vivo*. Our findings have important implications for the design and clinical translation of autologous and allogeneic organoid cellular therapies.

**ONE-SENTENCE SUMMARY:** The immune response to human primary cholangiocyte organoids is ameliorated by donor-recipient HLA matching.

## INTRODUCTION

Regenerative cellular therapies have emerged as a promising approach for the repair or replacement of diseased or damaged tissues and cells (*1*–*3*). The predominant strategy for the generation of therapies by differentiating embryonic stem (ES) or genetically-modified induced pluripotent (IPS) cells into the desired cell type (*1, 4*). Cells cultured in 3D organoid format have also recently been developed as an alternative strategy for the generation of cellular therapies (*5*–*7*). An important barrier for the clinical translation of cellular therapies is a thorough understanding of their immunogenicity (*8, 9*). This knowledge is critical for informing the use of the clinical strategies for the reduction of the immune response after transplantation, including the use of immunosuppressive drugs, donor-recipient matching, gene editing and encapsulation of the cells (*10*).

It is generally expected that ‘autologous’ cellular therapies, derived from cells obtained from the intended recipient, are unlikely to induce an immune response. However, early studies suggested that syngeneic (autologous) mouse iPSCs may or may not be immunogenic (*11*– *16*) and that immunogenicity of autologous iPSC-derived cellular therapies may be dependent on the target cell type and the immune microenvironment (*17*–*20*). These conflicting findings may be in part because generation of ES- and iPSC-derived cellular therapies necessarily involves genetic modification that may affect the immunogenicity of the derivative cells. Furthermore, ES- and IPS-derived cells may lack full differentiation into an adult phenotype, which may also impact their immunogenicity (*21*).

Development of autologous cellular therapies represents significant logistic and economic barriers that are likely to limit their widespread clinical use for the foreseeable future (*4, 22*). ‘Allogeneic’ cellular therapies, derived from a different donor than the intended recipient, are thus being explored as potentially cheaper ‘off the shelf’ treatments (*8*). Clinical data from decades of solid organ transplant experience demonstrate that the immune response to most allogeneic solid organs is driven predominantly by human leukocyte antigen (HLA) molecules. Consequently, when permitted by a large pool of donors and recipients (such as in kidney transplantation), HLA-matching between the donor and the recipient is an effective strategy for reducing (but not eliminating) the likelihood of immune-mediated rejection and for prolonging graft survival (*23*–*25*). Allogeneic cellular therapies are similarly expected to induce an immune response via direct and indirect allorecognition mechanisms driven in large part by HLA. However, it is important to note that most cellular therapies in advanced stages of development consist of a pure population of a single cell type (*26*–*28*), in stark contrast to solid organs that are composed of numerous cell types with diverse phenotypes and functions, including antigen presenting cells that are highly adapted to driving an immune response.

A key challenge to understanding the immune response to human cellular therapies has been the availability of appropriately refined experimental models and access to primary human tissues. Most studies to date have been performed exclusively *in vitro*, have used mouse-derived cellular therapies transplanted into wild-type immunocompetent mice, or used ES- or IPS-derived cells transplanted into mice reconstituted with an allogenic human immune system (*10*–*19*). Therefore, a detailed profiling of the human immune response under different HLA-matching scenarios using relevant humanized models and using cutting-edge technology is critical for the advancement of these therapies.

We have previously shown that primary cholangiocyte organoids (PCOs) can be derived from human cholangiocytes without genetic editing and have potential for use as cellular and bioengineered therapies for bile duct disorders (*29*–*31*). In this study, we use a comprehensive panel of *in vitro* and *in vivo* experimental models to examine, for the first time, the response of primary human immune cells to autologous and allogeneic human PCOs as an exemplar primary organoid cellular therapy. We demonstrate that autologous PCOs induce minimal immune response and that the immune response to allogeneic PCOs is driven by the level of HLA mismatch between the donor and recipient, showing evolving stages of immune rejection in a humanized mouse model. Our findings provide high-resolution analysis of immunogenicity of autologous and allogeneic organoid cellular therapies, highlighting important implications for their clinical translation.

## RESULTS

### PCOs upregulate expression of HLA I and HLA II in inflammatory conditions

Human PCOs were generated and characterised as described previously from gallbladder or bile duct biopsies (*29*–*31*) taken from more than 70 HLA-typed deceased transplant organ donors. For this work we selected one bile-duct derived PCO line which showed characteristic PCO spheroid morphology (**Fig S1A**), expression of specific cholangiocyte markers including *SOX17, SOX4, TFF2, FGF2, KRT-7* and *KRT-19* (**Fig S1B**), in addition to alkaline phosphatase (ALP) and gamma-glutamyl transferase (GGT) enzymatic activity comparable to primary cholangiocytes (**Fig S1C and S1D**).

Flow cytometric analysis revealed high bimodal expression of HLA-I and low expression (<40%) of HLA-II by primary cholangiocytes. PCOs cultured under normal conditions abundantly expressed HLA-I but no HLA-II. Both HLA-I and HLA-II were upregulated on PCOs when cultured in inflammatory conditions, simulated by 2 days of co-culture with IFN-γ. Primary cholangiocytes and PCOs all expressed the cholangiocyte marker CD326/Epcam as expected (**Fig. 1A**). The relative expression of HLA-I (A, B, C), and HLA-II (DP, DQ, DR) genes were quantified using qPCR (**Fig. 1B**), demonstrating upregulation by PCOs in inflammatory conditions for all loci and comparable to primary cholangiocytes. To examine whether the upregulation of HLA-II by PCOs upon co-culture with IFN-γ is physiologically relevant, we transplanted PCOs under the kidney capsule of mice reconstituted with an allogeneic human immune compartment (humanized mice). There was robust *in vivo* upregulation of expression of HLA-II by PCOs that were transplanted into humanized mice compared to immunodeficient mice (**Fig. 1C**). These data suggest that both HLA-I and HLA-II are likely to drive the immune response to PCOs in the clinical setting.

**FIG 1.**
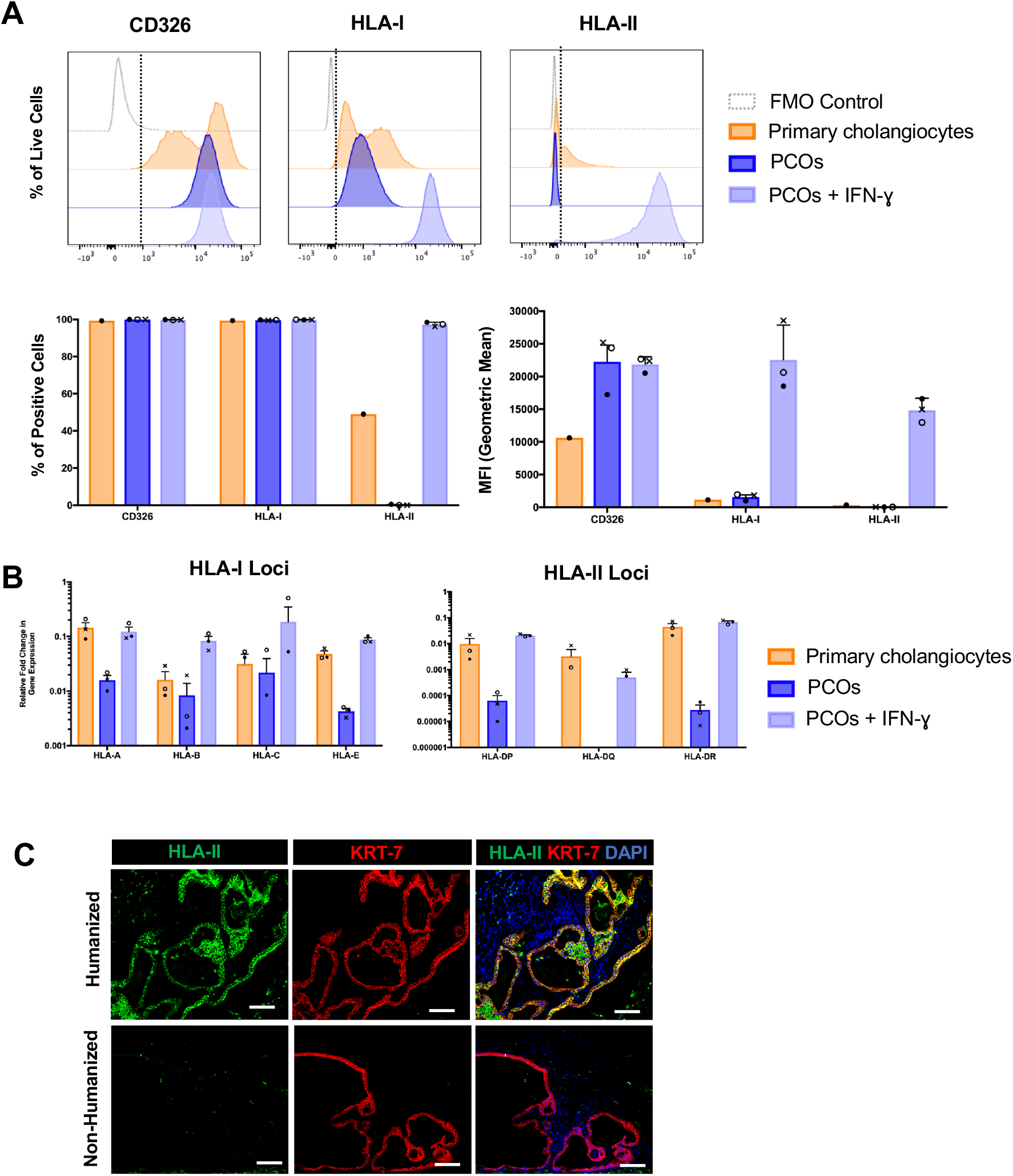
HLA expression by primary cholangiocytes and PCOs. (A) Representative flow cytometry charts (upper row), and bar graphs showing percentage and mean intensity fluorescence (MFI, lower raw) of expression CD326, HLA-I and HLA-II surface expression of primary cholangiocytes and PCOs cultured with or without IFN-γ stimulation for 2 days from three primary gallbladder/bile-duct derived lines (shown in different symbols). Fluorescence minus one (FMO) controls are used for gating. (B) qRT-PCR quantification of HLA-I (HLA-A, HLA-B, HLA-C, HLA-E) and HLA-II (HLA-DP, HLA-DQ, HLA-DR) expression by primary cholangiocytes and PCOs cultured with or without IFN-γ stimulation for 2 days from three primary gallbladder/bile-duct derived lines (shown in different symbols). Error bars represent mean±SEM from three PCO lines. (C) Immunofluorescence images showing expression of human HLA-II and KRT-7 in PCO grafts under the kidney capsule in immunodeficient NSG mice with and without reconstitution with allogeneic human immune cells (humanized). Error bars represent mean±SEM from three independent experiments. Scale-bars: 100µm.

### Donor-recipient HLA mismatch determines *in vitro* immune response to PCOs

We next assessed the level of immune activation as measured by cytokine secretion *in vitro* by co-culturing PCOs with spleen-derived mononuclear cells (SPMCs) derived from same deceased donors used to derive the PCO lines (**Table S1**), thus enabling immunological experiments with autologous or allogeneic donors with known HLA mismatch. In Autologous experiments, PCOs and SPMCs were obtained from the same deceased transplant organ donor. In Partial Match experiments, PCOs and SPMCs were matched at HLA-I only (A, B and C loci). In Full Mismatch experiments, PCOs and SPMCs were mismatched at both HLA-I and HLA-II loci (A, B, C, DP, DQ and DR) (**Table S2**). We used SPMCs (rather than peripheral blood mononuclear cells; PBMCs) for these experiments as it was possible to obtain much larger numbers of SPMCs per donor to enable both *in vitro* and *in vivo* experiments. Of note, we have previously shown that SPMCs can be used for immunological assays *in vitro* and *in vivo* (*32*). Fully mismatched PCOs induced the strongest immune response as measured by levels of IFN-γ, TNF-α and IL-6 secretion compared to other groups (**Figs. 2A-C. Table S3**). Except for IFN-γ, Autologous and Partial Match groups induced a similar level of immune activation that was higher than the negative controls; the level of IFN-γ production induced by Autologous PCOs was not different to negative controls. Secretion of the anti-inflammatory IL-10 was also significantly higher in the Full Mismatch group compared to other groups, potentially suggestive of simultaneous activation of inhibitory pathways (**Fig. 2D**). There were no changes in IL-12p70, IL-13, IL-1β, IL-2, IL-4 and IL-8 levels (**Table S3**).

**FIG 2.**
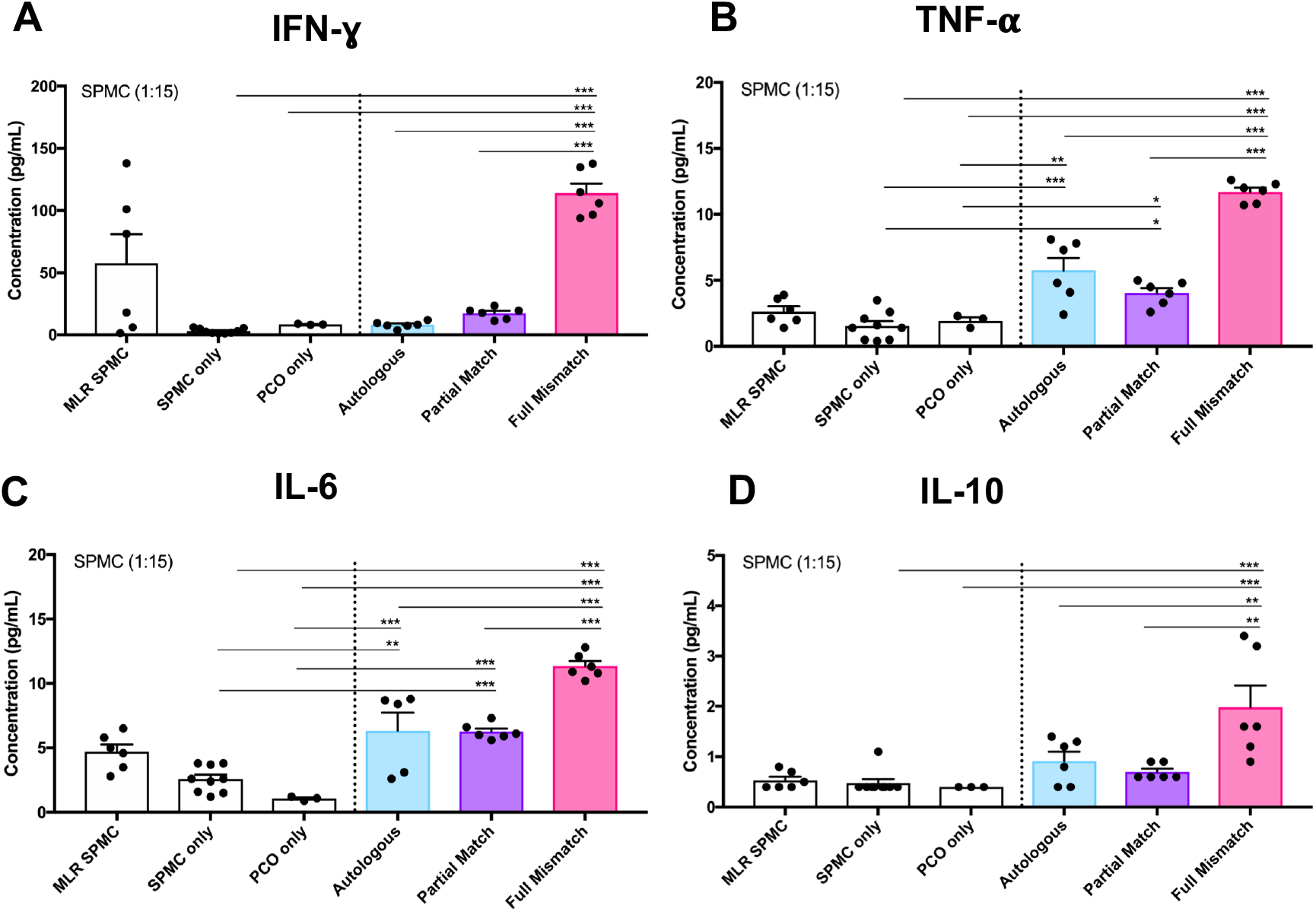
Activation of autologous and allogeneic lymphocytes by PCOs. (A) Bar graphs showing concentration of IFN-γ secretion by SPMCs when co-cultured (1:15 ratio) with Autologous, Partial Match and Full Mismatch PCOs for 5 days. Negative controls are PCOs only and SPMCs only and positive control is SPMC MLR from two different donors. Same conditions were analysed for TNF-α (B), IL-6 (C) and IL-10 (D). Error bars represent mean±SEM from three independent experiments. *p<0.0001; **p<0.001; *p<0.01.

### Magnitude of *in vivo* immune response to PCOs is driven by the level of HLA mismatch

We next examined the *in vivo* immune response to PCOs using a humanized mouse model. PCO fragments were injected under the kidney capsule of immunodeficient Nod Scid Gamma (NSG) mice. At 3 weeks post-injection, animals were humanized by intraperitoneal injection of Autologous, Partial Match or Full Mismatch SPMCs obtained from the same donors as used for the *in vitro* co-culture experiments. Importantly, in each animal, one kidney was transplanted with PCOs in Matrigel, and the contralateral kidney was injected with Matrigel only, thus serving as control for the effect of surgery and possible immune response to the Matrigel matrix. 6 weeks post-injection, animals were culled and their spleens were recovered to assess the human immune cell composition (**Fig. 3A**). Flow cytometry analysis demonstrated 20-40% human CD45-positive cells in the spleen, confirming successful humanization (**Figs. S2A-B**). CyTOF analysis revealed reconstitution with predominantly lymphoid cells as expected, with paucity of myeloid cells. Of note, the immune profile in the spleen of the mice was different in animals transplanted with Autologous, Partial Match or Full Mismatch PCOs (**Figs. 3B-D**). The Full Mismatch group had a decreased abundance of CD8 and a relatively higher proportion of B cells and CD57-Effector Memory CD4 T cells. Original donor SPMCs also showed differences in abundance of immune cell types (CD4, CD8, NK, Monocytes/DC, B cells), but with overall good representation of all cell types in all groups (**Fig. S2C**).

**FIG 3.**
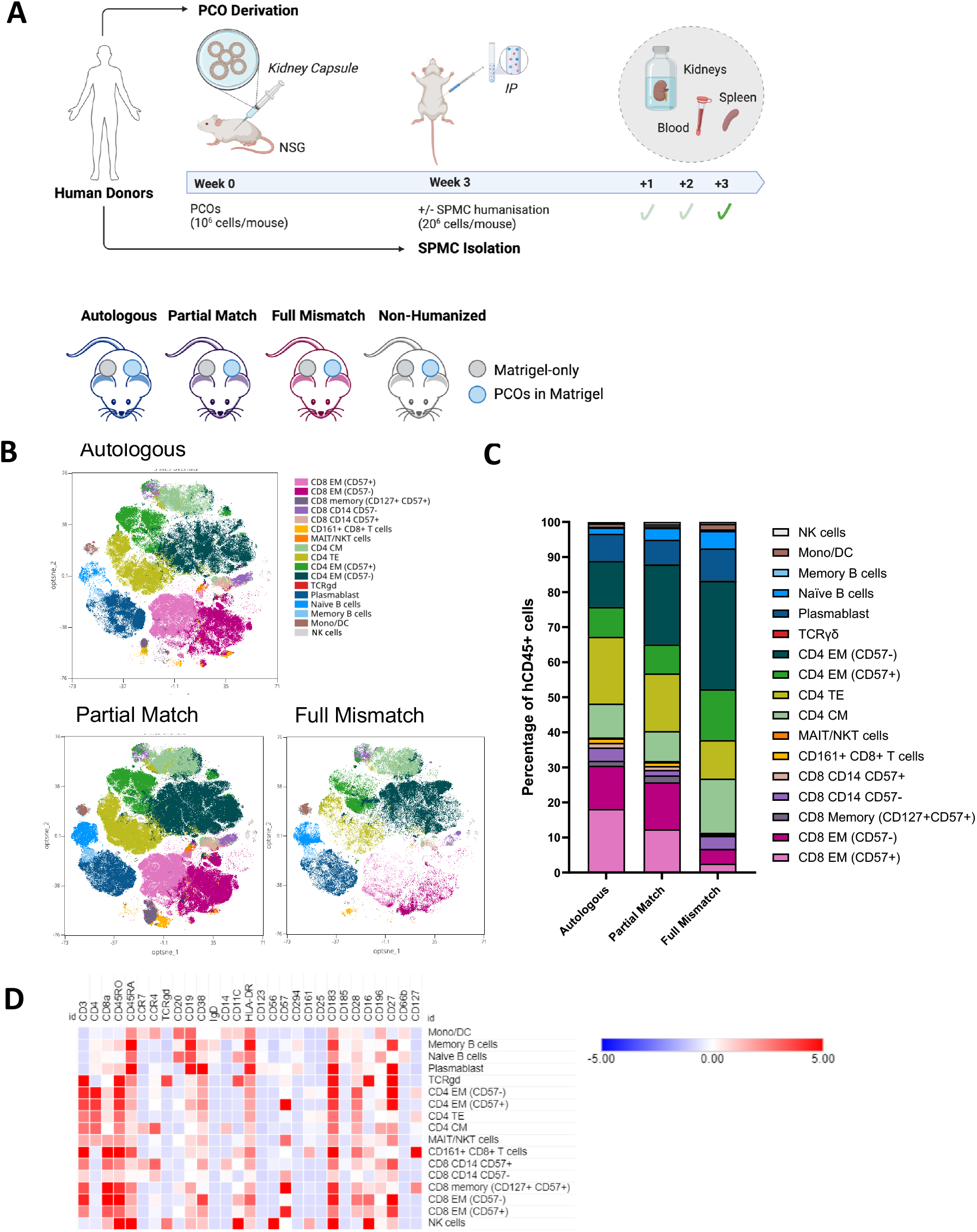
The immune profile of humanized mice transplanted with PCOs. (A) (Upper panel) Schematics of the PCO derivation, PCO injection and subsequent humanization with SPMCs after 3 weeks (n=5 per group). Animals were culled and samples (kidneys, spleens and blood) collected after 3 weeks of humanisation. (Lower panel) Representation of the groups included in the study: PCOs were injected in Matrigel under the left kidney capsule. Matrigel alone was injected under the contralateral kidney capsule to control for Matrigel-driven immune infiltration. Humanization was performed using Autologous, Partial match or Full mismatched human SPMCs. A non-humanized group was added as control. (B) OptSNE visualisation and FlowSOM clustering show the immune profile of human CD45+ cells with the distribution of immune cell subpopulations in different colours. Mice with the same donor engraftment have been overlaid. (C) Stacked bar graph representing the frequency of immune subsets in each type of engraftment. (D) Heatmap illustrating the level of expression of each marker for each cluster defined by FlowSOM analysis (all groups combined).

Assessment of local immune infiltration to the graft site was performed by immunofluorescence staining with specific human immune and cholangiocytes markers (CD45 and KRT7, respectively). PCO grafts had the characteristic morphology and surface marker expression in all groups (**Fig. 4A**). The entire engrafted areas were scanned, hCD45-positive cells were segmented and counted with a custom-made pipeline (**Fig. S3**). The ‘background’ hCD45+ immune infiltration in the matrigel-only area of the contra-lateral control kidney was then subtracted to correct for non-PCO-driven immune infiltration. The total PCO-specific immune infiltration was significantly greater in the Full Mismatch group, followed by Partial Match and Autologous groups (**Fig. 4B**). Importantly, Autologous PCOs induced an immune infiltration (∼5% of total cells) above that seen in the contralateral Matrigel-only control kidneys from the same mice. This apparent immune response to Autologous cells was consistent with possible low-level immune activation suggested by the *in vitro* co-cultures **(Fig. 2)**.

**FIG 4.**
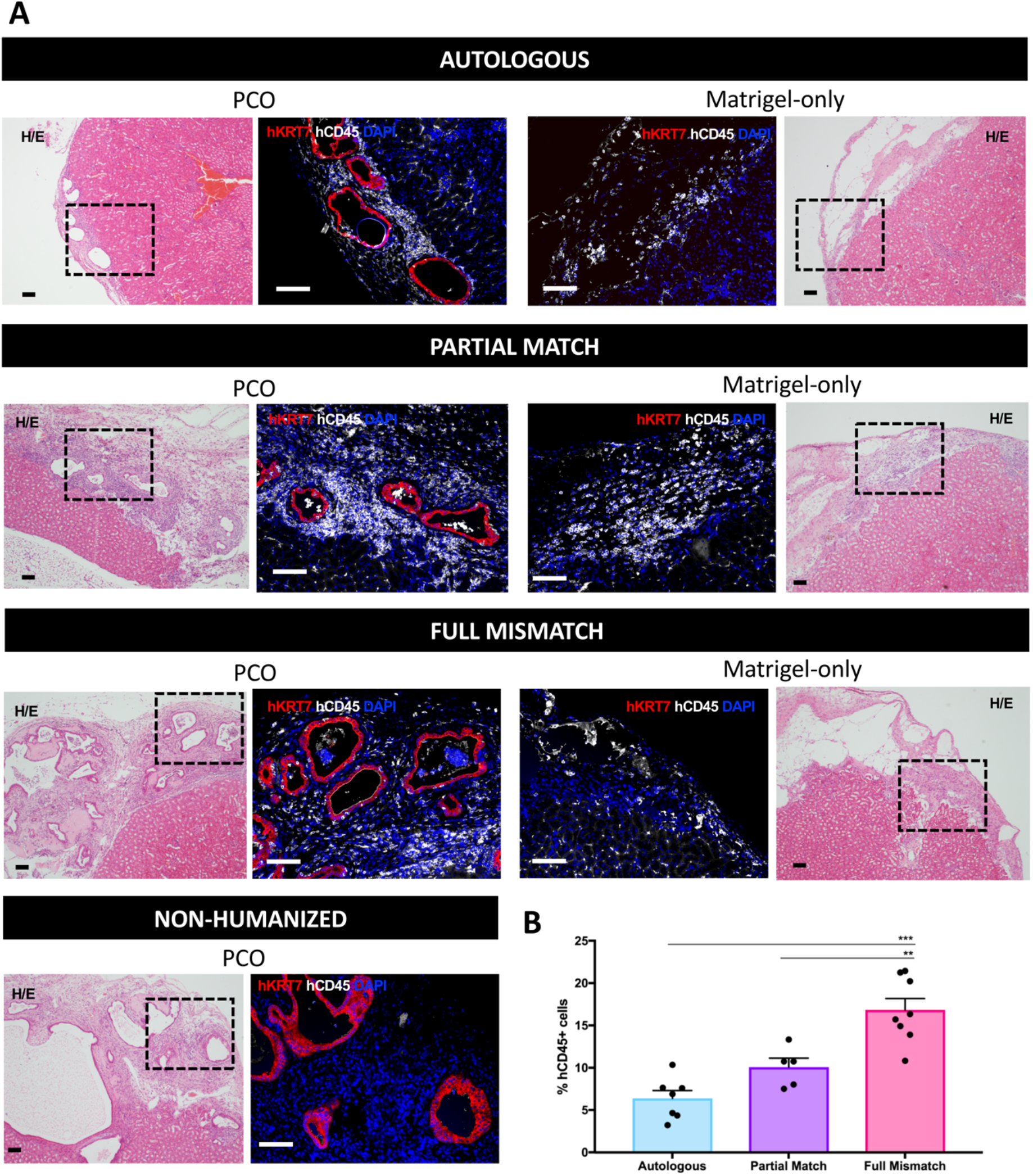
Infiltration of hCD45+ cells into PCO graft sites. (A) Hematoxylin/Eosin and immunofluorescence images of injected PCOs under the kidney capsule showing expression of human KRT-7 and human CD45 markers in Autologous, Partial Match, Full Mismatch and non-humanized groups. Matrigel-only images are shown as controls for background immune infiltration induced by surgical procedure and injection of Matrigel. Scale-bars: 100µm. (B) Percentage of hCD45 positive cells infiltrating into the graft site, above Matrigel-only controls, quantified in the respective groups. Error bars represent mean±SEM from 1-4 quantified areas per mouse per group. *p<0.0001.

### HLA mismatch determines phenotype of immune infiltration into the graft site *in vivo*

To determine the phenotype of the immune infiltrate into the graft site, we performed spatial transcriptomics using the Geomax platform (Nanostring), which enabled the retrieval of RNA from hCD45+ and KRT7+ cells. Deconvolution of the hCD45 compartment showed that the infiltrate in the Full Mismatch group had a decrease in CD4+ Memory T cells and an increase in Tregs, B cells (Naïve and Memory) and Monocytes relative to Matrigel-only controls (**Fig. 5A and Fig. S4A**). Difference in immune composition was less prominent between the Autologous and Partial Match groups, however, the former showed a shift from CD4 memory to CD8 memory; and the latter a change from CD4 native to CD8 memory. Overall, there was limited overlap in upregulated or downregulated genes between the groups and different immune-related pathways were enriched in the three groups (**Fig. 5B, Fig. S4B-C, Table S4)**. In particular, hCD45+ cells in the Autologous group did not show enrichment in any particular immune responses compared to controls (Matrigel-only). Conversely, the Partial Match group depicted general cellular immune activation, while the Full Mismatch group showed an antigen-driven, humoral response in addition to B cell and Treg pathways (**Fig. 5C**), which is consistent with the cell types described by the deconvolution. Interestingly, the KRT7+ compartment shared significantly more up- and down-regulated genes between groups (**Fig. S5A-B, Table S4**). Pathway enrichment analysis suggested that PCOs in the Partial Match group had upregulated pathways related to cell death state (e.g., apoptotic pathways), while Full Mismatch PCOs had upregulated cell stress (but not death; e.g., ubiquitin) pathways. Collectively, the data are consistent with the hypothesis that immune-related pathways are relatively quiescent in the Autologous group, but that there is ongoing activation of the immune pathways in the Partial Match and the Full Mismatch groups that have evolved to different stages of the immune response (**Fig. S5C**).

**FIG 5.**
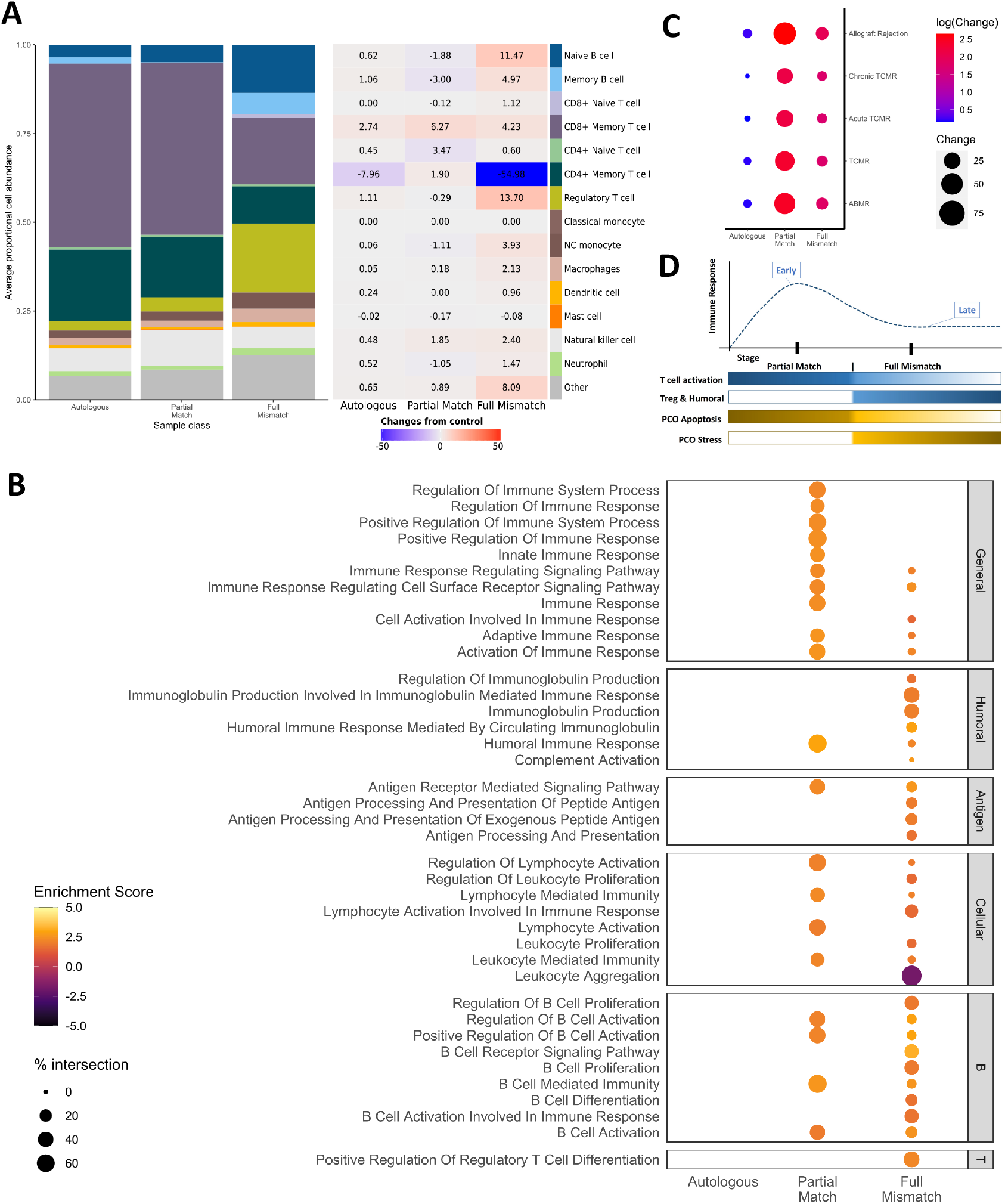
Phenotype of infiltrating immune cells into PCO graft sites. (A) Histogram showing abundance of different immune cells types infiltrating into PCO graft sites (left) and their respective fold change compared to Matrigel-only controls (right). Matrix (SafeTME) was extracted from Danaher et al (*58*). (B) Pathway enrichment analysis of Autologous, Partial Match and Full Mismatch groups for general, humoral, antigen, cellular, B cell and Treg pathways. (C) Dot plot showing the association of Autologous, Partial Match and Full Mismatch groups with rejection categories extracted from human kidney rejection datasets (TCMR, ABMR, chronic TCMR, acute TCMR, allograft rejection). (D) Schematic of the evolution of the stages of the human immune response to PCOs with different levels of matching in a humanized mouse model.

### Immune pathways activated by PCOs in humanized mice correlate with those activated by human organs undergoing rejection in transplant recipients

Our data suggest that, compared to the Partial Match group, the immune response in the Full Mismatch group is at a later stage of its evolution, with a dominance of B and regulatory T cells infiltrating the graft site and activation of the associated pathways. Conversely, Partial Match grafts appeared to have induced an earlier ‘active’ stage of the immune response. To examine this hypothesis, we assessed how the patterns of immune activation in the humanized mice compare to those seen in human transplant recipients of solid organs with biopsy-proven active clinical rejection. We thus compared the spatial transcriptomics data from the humanized mice to publicly available human datasets: two kidney rejection microarray datasets (*33, 34*); Banff-Human Organ Transplant (B-HOT) dataset (*35*) with classified T cell-mediated rejection (TCMR), antibody-mediated rejection (ABMR), Acute TCMR and Chronic TCMR; and allogeneic rejection pathways extracted from KEGG and PathCards (Allograft Rejection). Consistent with our hypothesis, we noted that the gene expression profile in human kidneys undergoing rejection correlated best with the Partial Match group, and less so with the Full Mismatch group (**Fig. 5C**). Overall, spatial transcriptomics in this SPMC-humanized mouse model captured at a single snapshot in time (6 weeks post-cell therapy injection), suggest that the immune response elicited by Partial Match PCOs corresponds to early stages of response with potent T cell activation and with PCOs showing transcriptional signs of apoptosis. Conversely, Full Mismatch PCOs have induced an immune response that has evolved to a later stage involving B cells, Tregs and humoral response (**Fig. 5D**).

## DISCUSSION

To our knowledge, this is the first comprehensive study comparing the *in vitro* and *in vivo* immune responses to autologous and allogeneic healthy organoids derived from primary cells. We selected PCOs as our exemplar genetically-unedited organoid cellular therapy, as they have previously been shown to have potential utility as a cellular therapy (*29*–*31*). We show that human PCOs upregulate HLA-I and HLA-II molecules in inflammatory conditions *in vitro* and *in vivo*. The immunogenicity of PCOs *in vitro*, as quantified by release of inflammatory cytokines by co-cultured lymphocytes, was dependent on the level of donor-recipient HLA matching. Consistent with this finding, immune infiltration into the graft site in humanized mice was also dependent on the level of donor-recipient HLA matching. Autologous PCOs also induced low-level immune infiltration into the graft site compared to controls. Spatial transcriptomics data indicate evolving stages of immune rejection dependent on donor-recipient HLA matching. Specifically, the immune infiltrate profile in the Partial Match graft sites was consistent with an early and prominent T cell-driven response, compared to a later stage immune response driven by B cells and Tregs in the Full Mismatch group. Finally, we were able to correlate the immune profile of the graft-infiltrating cells onto previously published human kidney allograft rejection profiles.

Here we focused on the impact of HLA on immunogenicity of the PCOs, because decades of evidence from solid organ transplantation has established HLA as the predominant driver of the alloimmune response and demonstrated the beneficial impact of donor-recipient HLA matching on graft immune rejection and survival (*23*–*25*). To enable this study, we obtained paired primary immune cells and primary cholangiocytes from a large number of deceased transplant organ donors. By obtaining large numbers of spleen-derived lymphocytes from every donor, we were able to replicate the same experiments *in vitro* and *in vivo*. Similarly, by generating large number of PCO lines, we were able to compare the immune response to Autologous PCOs, PCOs matched at HLA-I (A, B and C) but mismatched at HLA-II (DP, DQ and DR) loci (Partial Match group) and PCOs mismatched at both HLA-I and HLA-II loci (A, B, C, DP, DQ and DR).

Our data demonstrate that PCOs, similar to primary cholangiocytes, upregulate both HLA-I (A, B and C) and HLA-II (DP, DQ and DR) in inflammatory conditions *in vitro* and *in vivo*. Consistent with our findings, upregulation of HLA in inflammatory *in vitro* conditions has been previously reported for other putative cellular therapies derived from iPS or ES cells (*36*–*39*). The magnitude of the *in vitro* and *in vivo* immune responses (as measured by the total number of CD45+ infiltrating immune cells) to allogeneic PCOs was significantly reduced by donor-recipient matching at HLA-I but was nonetheless greater than autologous control PCOs, driven by mismatch at HLA-II. These findings are indicative that allogeneic PCOs would induce both anti-HLA-I and anti-HLA-II immune responses in a clinical setting, and that donor-recipient HLA matching, at both HLA-I and HLA-II loci, would be an effective strategy for amelioration of the alloimmune response in patients.

The immune profile of the lymphocytes infiltrating into the graft site *in vivo* was different depending on the level of HLA mismatch. Spatial transcriptomics analysis revealed differences in immune cell populations and cell enrichment pathways present at the graft site depending on the level of HLA mismatch. The immune infiltrate in the Partial Match group was dominated by CD8+ and CD4+ T cells, but with B cells and Tregs in the Full Mismatch group. These findings suggest the dominance of two different phases of immune response, possibly corresponding to an early/prominent vs a late/resolved response, respectively. The prominent immune profile captured in the Partial Match group was highly associated with human rejection signatures extracted from datasets of human kidneys. Although PCO grafts demonstrated characteristic morphology when stained by specific cholangiocyte markers for all groups, spatial transcriptomics suggested a more apoptotic state in the Partial Match group. These results are consistent with an evolving immune response, comprising a more vigorous early immune response captured in the Partial Match group, and a later quiescent stage in the Full Mismatch group. Interestingly, we also observed greater production of IL-10, a predominantly anti-inflammatory cytokine, in lymphocyte co-cultures with Full Mismatch PCOs. Release of IL-10 and other inhibitory cytokines has indeed been postulated as one of the negative feedback mechanisms to curtail ongoing immune responses to alloantigens, including from NK cells (*40*). However, we were unable to experimentally test how donor-recipient HLA mismatch influences the kinetics of the evolution of the immune response to PCOs, because all animals were culled at a single fixed time point after immune reconstitution. It is also possible that differences in the donor immune compartment or immune engraftment in the animals contributed to some of the differences in the immune responses captured in our study. Of note, however, the composition of the immune compartment of the donors were broadly similar.

The immune response to autologous human cellular therapies have been reported in previous studies with conflicting findings (*17*–*19, 41*–*43*). Our data suggest that autologous PCOs may also induce a low-level immune response. Autologous PCOs induced *in vitro* production of higher levels of the inflammatory cytokines TNF-α and IL-6, but not IFN-γ. Autologous PCOs also induced greater infiltration of immune cells into the graft site compared to control injection sites. By injecting Matrigel alone under the contralateral kidney capsule in every humanized mouse, we controlled for both the immune response induced by surgery and Matrigel itself, as well as for the inter-animal variance in immune engraftment that is inevitable with humanised mouse models (*44*–*46*). However, it is not possible to definitively conclude whether autologous PCOs would induce a similar immune response in a clinical setting, or if such weak immunogenicity would be clinically relevant and lead to significant cell loss or require short-term immunosuppression.

There are several possible explanations why autologous PCOs may have induced an apparent low-level immune response in our models. Some cell death is inevitable when cells are cultured *in vitro* and after manipulation for transplantation *in vivo*, which can induce an immune response (*47, 48*). The apparent low level immune response to autologous PCOs may, therefore, represent a self-limiting non-specific immune activation that is of minimal clinical significance. Our data confirm that culture conditions affect expression of HLA on PCOs; it is thus possible that other as-yet-unidentified self-antigens are also up- or down-regulated in culture, potentially inducing a weak immune response. Moreover, Matrigel is a mouse-derived hydrogel (*49*) and the presence of xeno-antigens in culture could have affected the immunogenicity of PCOs, especially since they express the cellular machinery for exogenous antigen presentation (HLA-II). An immune response driven by xeno-antigens, if indeed present, would be expected to be eliminated if PCOs are cultured under GMP conditions for clinical use. Finally, although our PCOs were not genetically edited, it is well established that cells cultured *in vitro* can acquire mutations with passaging (*50, 51*), as is also the case with all human cells that divide during embryonic development and ageing in the donor (*52, 53*). It is, therefore, possible that expression of unidentified neoantigens can derive an immune response in autologous cells.

In our in vivo experiments, we transplanted PCOs under the kidney capsule rather than into the liver. We chose the kidney capsule because it is a highly vascular niche (*54, 55*) that supports the survival of PCOs, enables localization of the graft at the experimental endpoint, and represents a well-characterised niche for assessment of immunogenicity. It is possible, however, that the immune response to PCOs may differ qualitatively or quantitatively from our findings when transplanted into the liver in patients. While we used unselected SPMCs (consisting of both lymphoid and myeloid cells) in our *in vitro* experiments, our humanised mouse models primarily recapitulated the lymphoid human immune compartment. It is thus possible that recipient myeloid cells may also influence the immune response to PCOs in a clinical setting. Future studies utilizing other experimental models would be necessary to examine the contribution of myeloid cells to the alloimmune response to PCOs. Notwithstanding these caveats, it is important to note that our *in vitro* and *in vivo* experimental models were sufficiently refined to distinguish between the immune response to allogeneic PCOs with different levels of donor-recipient HLA mismatch.

In summary, our study confirms that both HLA-I and HLA-II are likely to drive the immune response to organoid cellular therapies in a clinical setting. Donor-recipient HLA-matching, and elimination or HLA expression by genetic editing, are likely to be effective strategies to substantially ameliorate, but not eliminate, the immune response to allogeneic cellular therapies in a clinical setting. This is consistent with transplant studies that have identified minor histocompatibility (and other) molecules as making a small contribution to the alloimmune response (*56*). Our findings suggest that some immunosuppressive therapies are still likely to be required in patients receiving optimally-HLA-matched cellular therapies. Moreover, our data also do not exclude the possibility that autologous cellular therapies may similarly induce a low-level immune response in patients. However, it is well established that the adverse effects of immunosuppressive therapies decrease with reduced total burden of immunosuppression (*57*). Elimination of the alloimmune response against HLA-I and HLA-II, through the use of autologous cellular therapies, HLA-matching or genetic editing, would thus be expected to substantially reduce the required dose of immunosuppression and associated adverse effects in patients. Taken together, this work provides a high-resolution perspective to guide future efforts focused on deeper understanding of the immunogenicity of cellular therapies, and accelerate the clinical translation of cell-based products for regenerative medicine applications.

## ACKNOWLEDGEMENTS

We are grateful to the donors and their families and the Cambridge Biorepository for Translational Research (CBTM) for the previous gifts of tissue donation. We would also like to thank Jasper Callemeyn and Marteen Naesens for their input on the analysis of human kidney transplantation datasets and Fotios Sampaziotis for his input into the generation of cholangiocyte organoids. Figure drawings were created with BioRender.com.

## Funding

S.P.-R. was supported by awards from Medical Research Council (MRC) UK Regenerative Medicine Platform (MR/S020934/1) and MRC Confidence in Concept (G116517). O.T was supported by an award from the MRC. L.V. was supported by European Research Council Grant New-Chol (ERC: 741707) award.

## Author Contributions

S.P.-R. conceived and designed the study; performed experiments; acquired, interpreted, and analysed the data; drafted and edited the manuscript. O.T. conceived and designed the study; generated PCO lines and performed experiments. W.L. performed bioinformatic analysis. M.M. and C.B. performed Cytof experiments and analysed data. T.T. and S.R. contributed to tissue sectioning experiments. K.M. provided primary tissue and isolated splenocytes. J.J and C.B performed slide scanning and hCD45 quantification. N. H. provided funding and supported data analysis. L.V. provided funding and supported generation of primary cholangiocytes organoids. K.S.-P. Conceived and designed the study, performed animal experiments, interpreted data, wrote and edited the manuscript. All authors approved the manuscript.

## Competing Interests

K.S.-P. and L.V. are founders and shareholders of Bilitech Ltd and inventors on the patent applications GB/19.96.17 and GBA 201709704 held by the University of Cambridge that cover the derivation and use of cholangiocyte organoids for regenerative medicine. The remaining authors have no competing interests to disclose.

## Data and Materials Availability

All data are available in the main text or the supplementary materials. Spatial transcriptomics data (processed count matrices, annotation files, meta data) are available on Dryad (DOI:10.5061/dryad.g4f4qrfx9). Code for analyses are shared at Zenodo (DOI:10.5281/zenodo.10054742). Cholangiocyte organoids described in this publication are available from University of Cambridge under a material transfer agreement with the university.

## Notes

### Competing Interest Statement

The authors have declared no competing interest.

